# Population scale nucleic acid delivery to *Caenorhabditis elegans* via electroporation

**DOI:** 10.1101/2020.10.15.340513

**Authors:** Anastasia S. Khodakova, Daniela Vidal Vilchis, Ferdinand Amanor, Buck S. Samuel

## Abstract

The free-living nematode *C.elegans* remains one of the most robust and flexible genetic systems for inter-rogating the complexities of animal biology. Targeted genetic manipulations, such as RNA interference (RNAi), CRISPR/Cas9- or array-based transgenesis, all depend on initial delivery of nucleic acids. Delivery of dsRNA by feeding can be effective, but expression in *E. coli* is not conducive to experiments intended to remain sterile or with defined microbial communities. Soaking-based delivery requires prolonged exposure of animals to high material concentrations without a food source and is of limited throughput. Last, microinjection of individual animals can precisely deliver materials to animals’ germlines, but is limited by the need to target and inject each animal one-by-one. Thus, we sought to address some of these challenges in nucleic acid delivery by developing a population-scale delivery method. We demonstrate efficient electroporation-mediated delivery of dsRNA throughout the worm and effective RNAi-based silencing, including in the germline. Finally, we show that guide RNA delivered by electroporation can be utilized by transgenic Cas9 expressing worms for population-scale genetic targeting. Together, these methods expand the scale and scope of genetic methodologies that can be applied to the *C.elegans* system.

## INTRODUCTION

Understanding gene function is an essential task of modern biology. The nematode *Caenorhabditis elegans* is one of the most widely used and versatile animal models for studying nearly all aspects of animal biology [Corsi et al. 2015, Meneely et al. 2019]. For many years *C.elegans* has proven to be an effective and powerful genetically tractable system for functional characterization of genes in a whole organismal context [Housden et al. 2017]. First, *C.elegans* allows for a rapid analysis of gene function carried out via targeted RNA interference (RNAi)-based knock-down of gene expression [Timmons and Fire 1998, Conte et al. 2015]. Second, transgenic animals bearing exogenous genes can be created via microinjection of DNA constructs into the animal’s gonad resulting in the formation of heritable extrachromosomal arrays [Berkowitz et al. 2008]. Third, CRISPR/Cas9 genome editing tools have been developed for precise genomic manipulations that allow desired *C.elegans* mutants to be engineered [Dickinson and Goldstein 2016]. One of the critical steps for every genome manipulation pipeline is the delivery of nucleic acids inside the cell or animal. For *C.elegans*, microinjection of individual worms is a crucial step in the delivery of exogenous material. Microinjection remains the most time- and labor-intensive procedure for most *C.elegans* laboratories, whereas many other methods and approaches have been developed for different cellular and organismal systems [Alsaggar and Liu 2015]. Among others, electroporation has been recognized as a powerful and quick method for simultaneous nucleic acid transfer in large populations of bacterial, yeast and mammalian cells [Young and Dean 2015].The electric pulse applied to the cell destabilizes its membrane and causes formation of transient pores allowing exogenous material such as DNA, RNA, and proteins to enter the cell. Electroporation can also be used for introduction of exogenous material into entire tissues of the whole organism - e.g., electroporation of DNA in zebrafish [Kera et al. 2010], Xenopus [Haas et al. 2002], or silkworms [Ando and Fujiwara 2013]. However, this delivery method has not yet been applied to *C.elegans* animals. In this study, we demonstrate the feasibility and potential of the electroporation-based delivery of nucleic acids in *C.elegans* at a population scale. We show that electroporation-based delivery of double stranded RNA (dsRNA) triggers RNAi gene silencing pathways inside *C.elegans*. This protocol is accomplished at the scale of hundreds of animals, making it broadly applicable and useful for nucleic acids delivery. Finally, we show in proof-of-principle studies that electroporation-mediated delivery of single-stranded guide RNA (gRNA) molecules can be utilized to disrupt genes in the progeny of Cas9 expressing animals. Together, we anticipate electroporation-based methods to greatly enhance the scope and scale of genetic targeting in this already robust genetic system.

## MATERIALS AND METHODS

### Worm strains and maintenance

All strains were cultured on Nematode Growth Medium (NGM) plates seeded with *Escherichia coli* strain OP50 at 20°C. Mutant strains VC1119 *[dyf-2&ZK520.2(gk505) III]* (referred as *[sid-2(gk505) III]* in current study) and HC196 *[sid-1(qt9) V]* were obtained from the Caenorhabditis Genetic Center. Transgenic GR1403 *[Is(sur-5::gfp) I; eri-1(mg366) IV]* strain was a kind gift from Gary Ruvkun. The BIG0105 *[Is(sur-5::gfp) I]* strain was produced by crossing GR1403 with the Samuel lab stock of N2. Strains BIG0106 *[sid-1(qt9) V; Is(sur-5::gfp) I]* and BIG0107 *[sid-2(gk505) III; Is(sur-5::gfp) I]* were generated by crossing of HC196 and VC1119 mutants with BIG0105 strain. Transgenic strain EG9888 that stably expresses Cas9 in the germlines of animals was kindly gifted by Dr. Matthew Schwartz and Dr. Erik Jorgensen [*W01A8.6(oxTi—[Pmex-5::cas9(+ smu-2 introns), Phsp-16.41::Cre, Pmyo-2::2xNLS-CyOFP + lox2272])I*]. A complete list of worm strains used and prepared in this study can be found in **Table S1**.

### Synchronization

Nematodes were synchronized by bleaching and allowed to hatch overnight in M9 buffer [Stiernagle 2006]. Density of synchronized L1 larvae population was then measured.

### Production of dsRNA

PCR products corresponding to *gfp*, *dpy-13*, *nhr-23* and *pos-1* genes were generated with T7 primer (5’-AATACGACTCACTATAG-3’) and vectors isolated from the RNAi *E.coli* clones, using the following cycling conditions: 98°C 15 sec, 55°C 15 sec, 72°C 60 sec for 30 cycles. PCR product purification was performed according to manufacturer’s protocol with QIAquick PCR Purification Kit (Qiagen). Purified PCR products were then used as templates for in vitro transcription per AmpliScribe T7 High Yield Transcription Kit (Epicentre Technologies) specifications to obtain dsRNAs.

### Production of guide RNA

Production of the short gRNA (100 nt in length) specific to *dpy-10* gene was performed according to the protocol described in [Hwang et al. 2013]. In brief, a plasmid encoding gRNA (targeting *dpy-10*) was constructed as follows: pDR274 vector [Addgene plasmid 42250] for in vitro gRNA production containing a T7 promoter upstream of gRNA scaffold sequence was digested with BsaI enzyme (NEB). It was then used as a backbone for cloning the annealed oligonucleotides (dpy-10T: 5’-TAGGGCTACCATAGGCACCACGAG-3’; dpy-10B: 5’-AAACCTCGTGGTGCCTATGGTAGC-3’), containing *dpy-10* protospacer sequence (5’-GCTCGTGGTGCCTATGGTAG-3’). The sequence verified expression vector was then digested with HindIII enzyme (NEB) and used as a template for in-vitro transcription of gRNA by AmpliScribe T7 High Yield Transcription Kit (Epicentre Technologies).

### Electroporation of L1 worms with dsRNA

An aliquot of the synchronized worms of measured density was spun down at 500 rcf for 2 min to provide approximately 250 worms (unless otherwise specified) in volume of 5 *μ*L after the centrifugation. Then 5 *μ*L of worms were mixed with 40 *μ*L of electroporation buffer (Gene Pulser Electroporation buffer, Biorad) in 1.5 mL tubes, and allowed to incubate on ice for 5 min. An aliquot of 5 *μ*L of purified dsRNA (10 *μ*g/*μ*L) was added to the worms just before the electroporation, mixed by pipetting, and transferred to 0.2 cm electroporation cuvettes (Biorad). Animals were electroporated at 300 V for 10 ms (unless otherwise specified) by square-wave single pulse using a Bio-Rad Gene Pulser (BioRad). Immediately after the electroporation, worms were washed with 1 mL of pre-chilled M9 buffer, transferred into 1.5 mL tubes and centrifuged for 2 min at 500 rcf. Supernatants were discarded and animals were then transferred to *E.coli* OP50 seeded plates and cultured for 48 hours at 20°C.

### Electroporation of L4 and Young Adult animals

Synchronized L1 larvae worms were cultured on OP50 plates until L4 (55 hrs) or Young Adult (70 hrs) stage at 20°C. Then worms were washed off the plate with M9 buffer, followed by two additional washes in the same buffer to eliminate bacteria. Electroporation procedure for L4/YA worms was performed the same way as described for L1 worms. After the electroporation worms were allowed to recover and lay eggs on OP50 plates for 24 hours and then removed. Progeny development was monitored for 48 hours (unless otherwise specified) and worms were imaged.

### Image acquisition and analysis

Microscopy-based analyses were used to count animals, measure body size and GFP fluorescence intensity. For imaging, worms were washed off the OP50 lawn with M9 buffer containing 20 mM of NaN_3_, washed with the same buffer two times to remove bacteria and then transferred to wells of a 96-well plate or glass slide with a 2% agarose pad. Animals were imaged using the Eclipse Ti-5 fluorescence microscope (Nikon) with 4*×* and 10*×* and 20*×* magnification under non-saturating conditions. Analysis of imaging data was performed using Fiji software [Schindelin et al. 2012] and custom written MATLAB (Mathworks) scripts (**File S1**). A minimum of 50 animals were analyzed per group for worm body length measurement and GFP fluorescence (unless otherwise specified). Worm GFP fluorescence were calculated by dividing the sum of GFP intensities of all pixels over the total pixel number for each worm. Then the background fluorescence, calculated as average fluorescence intensity of all pixels in a region without worm, was subtracted from worm fluorescence. GFP fluorescence per worm is defined in arbitrary units (a.u.). Time-lapse bright field images of live worms with Dumpy and Roller phenotypes were used to create .mp4 video files of the worm’s movement (**File S2**).

### Genotyping and Illumina sequencing

Genotyping of generated BIG0106 *[sid-1(qt9) V; Is(sur-5::gfp) I]* and BIG0107 *[sid-2(gk505) III; Is(sur-5::gfp) I]* transgenic strains was performed using single worms PCR and primers listed in **Table S2** followed by Sanger sequencing confirmation of generated PCR products. In order to identify presence of CRISPR editing in *dpy-10* gene after the gRNA electroporation, single worm PCR products were analyzed by Illumina sequencing using 2*×*250 bp pair-end run. Primers were designed to generate 450 bp PCR product with gRNA target sequence located in the middle of the amplicon. Worms were lysed in DNA Quick Lysate (Epicentre Technologies) for 1 hour at 60°C and the lysate was then used as a template for PCR with Q5 Hot Start High-Fidelity DNA Polymerase (NEB). PCR products were purified using QIAquick PCR Purification Kit (Qiagen). Barcoded libraries production and Illumina sequencing run were provided by GENEWIZ. Two FASTQ files (R1 and R2) were generated for each sample (**File S3**), and subsequently analyzed using Cas-Analyzer online tool [Park 2017].

### Statistical analysis

Comparison of multiple groups was performed using the analysis of variance (ANOVA) with Bonferroni correction. P - values *<* 0.05 were considered statistically significant. All experiments were performed at least two independent times.

### Data availability

All *C. elegans* strains, primers and plasmids described in this study are available upon request. Raw data, scripts used for analyses and sequencing datasets can be found in supplementary files deposited on Figshare.

## RESULTS

### Initial considerations in development of an electroporation pipeline for *C.elegans*

Based on applications in other systems, we first established a reliable and robust pipeline for electroporation of *C.elegans* (outlined in **Figure 1a**) that could serve as a basis for further optimization. Briefly, one part of the worm suspension with desired number of worms is mixed with one part of the nucleic acid solution and eight parts of the electroporation buffer on ice to preserve the integrity of nucleic acids. The mixture then is transferred into the cuvette and electroporated under desired conditions. In current study we electroporated worms in 50 *μ*L of final solution. In total, the electroporation procedure was rapid (15 min) and tolerated by the animals.

**Figure 1:**
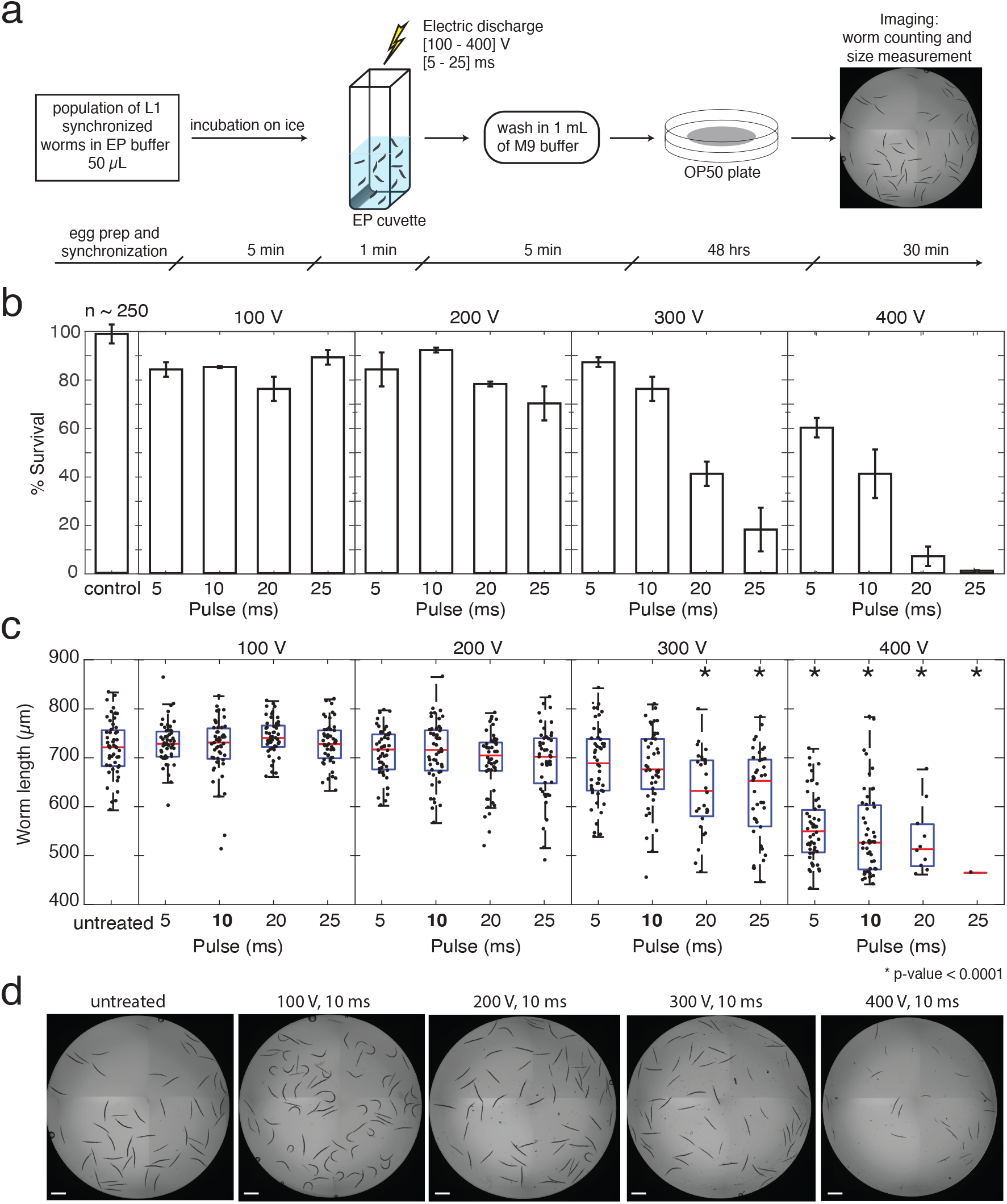
Optimization of electroporation conditions for *C.elegans* viability. General pipeline **(a)** of the electroporation procedure starts with the preparation of L1 synchronized worms (*~*250), which are then mixed with electroporation buffer 80% in chilled cuvettes. After electroporation, worms are washed with 1 mL of M9 buffer and collected by centrifugation at 500 rcf for 2 min, then transferred to *E.coli* OP50 seeded plates to grow for 48 hours at 20°C. Animal survival rates **(b)** and body lengths **(c)** varied based on the electroporation conditions applied. The evaluation was performed using N2 animals for each pair of electroporation parameters with an electroporation pulse duration ranging from 5 to 25 ms and voltage ranging from 100 to 400 V. Animals placed in electroporation buffer without electric discharge were used as “untreated” controls. Worm survival rates: mean *±* SD (standard deviation)% of two independent experiments. Body length measurements: red lines indicate means, blue boxes show 25^*th*^ and 75^*th*^ percentiles, whiskers show the data distribution range. *P - values *<* 0.05 were considered statistically significant (ANOVA test with Bonferroni correction). Representative images **(d)** of worm populations exposed to electric discharges of different voltages (10 ms pulses) demonstrate the pronounced effect of the electroporation procedure on animal viability. Scale bar = 500 *μ*m.

### Optimization of electroporation conditions for nucleic acids delivery while preserving animal viability

The efficiency of *in vivo* electroporation as a delivery tool is represented by an intersection of two key metrics: (1) maximum viability of worms under applied electroporation conditions and (2) the degree of material delivery itself. During electroporation, an electrical pulse is applied across the animal’s body with the assumption that some tissues may be more impacted than others. The cuticle, a multi-layered collagen outer tissue akin to our skin, provides considerable protection for the worm’s body and is likely to be a strong barrier for the electric pulses to bridge. To address these challenges, we sought to identify optimal electroporation parameters, in particular - pulse voltage and pulse length, that minimize adverse effects on worm physiology and maximize potential for nucleic acid delivery. These parameters were tested pairwise across a range of conditions for their impact on survival and developmental rates on populations of L1 synchronized N2 animals (*~*250). Microscopy-mediated worms’ assessment was performed after 48 hrs of recovery on *E. coli* lawns. Robust animal viability was observed (*>* 70%) at lower voltages (100-200 V) regardless of the pulse duration, and up to 10 ms pulses for 300 V treatments (**Figure 1b**). Beyond these conditions, treatment of worms at or above 300 V for longer than 20 ms significantly decreased animal survival rates (**Figure 1b**). Based on measurements of animal length and vulval morphology, similar combinations of high voltage and long duration of the electric pulse caused significant developmental delays in electroporated worms compared to the untreated control animals (**Figure 1c-d**). Fecundity rates of electroporated L1 worm populations under favorable conditions (at or below 300 V and 10 ms) also appeared to be similar to untreated controls (data not shown). Thus, treatment of worms with 300 V for pulse durations up to 10 ms minimizes adverse effects on animal viability and developmental timing while maximizes potential for material delivery. As we were also interested in the delivery of nucleic acids to the germlines, we performed similar optimization of electroporation parameters on L4 animals (N2) that are closer to reproductive maturity. Using the same matrix of voltage and pulse duration times as for L1 animals before, favorable viabilities remained *>*70% up to 400 V 10 ms (**Table S3**). Despite the apparent resilience in L4 animals from a viability perspective, we observed a collapse of one or more of the gonads in up to *~*15% of the cases starting from 300 V and 20 ms and onward, and as a consequence, a decrease in fecundity rate (data not shown).

### Evaluation of the effectiveness of electroporation of dsRNA in *C.elegans* populations

Silencing by RNAi in *C.elegans* is a sensitive method for specific knockdown of gene expression [Conte et al. 2015], and when applied to fluorescent transgenes, RNAi provides a robust visual phenotypic readout of the degree of knockdown at a cellular level. In *C.elegans*, RNAi-mediated silencing can be achieved by feeding worms with *E.coli* expressing a gene-specific dsRNA [Timmons and Fire 1998] or via soaking of worms in a highly concentrated solution of dsRNA ranging from 0.5-5 *μ*g/*μ*L [Ahringer 2006]. Ingested dsRNAs are recognized by the lumenal receptor SID-2 and subsequently engulfed [McEwan et al. 2012]. Engulfed dsRNAs are released and spread into the cell cytosol (and throughout the animal) via SID-1 membrane channels [Wang and Hunter 2017]. The presence of dsRNA in the cytosol triggers canonical RNA dependent RNA polymerase (RDRP)-based amplification and ultimately RNAi silencing of target genes [Shih and Hunter 2011]. In order to test the effectiveness of electroporation, we utilized this highly sensitive system to identify animals and tissues that were effectively delivered dsRNAs. To do this, we used transgenic animals BIG0107 that both produce GFP ubiquitously in the nuclei of all somatic cells and lack the ability to take up dsRNA from the intestine. Synchronized L1 populations of animals were electroporated using favorable conditions identified above and monitored for *gfp* silencing as a proxy for effectiveness of dsRNA delivery. Though all treatments with 100 V did not result in silencing, we observed significant reductions in GFP fluorescence in animals treated with 200 V or greater compared to controls (**Figure 2a**). Based on phenotypic analyses of the electroporated animals, we identified that treatments of animals with 300 V for 10 ms yielded the highest percentage of animals in the completely silenced (all but neuronal cells) category at nearly 60% (**Figure 2b**). Together, these results identified effective conditions that allow the delivery of dsRNAs into *C.elegans* animals.

**Figure 2:**
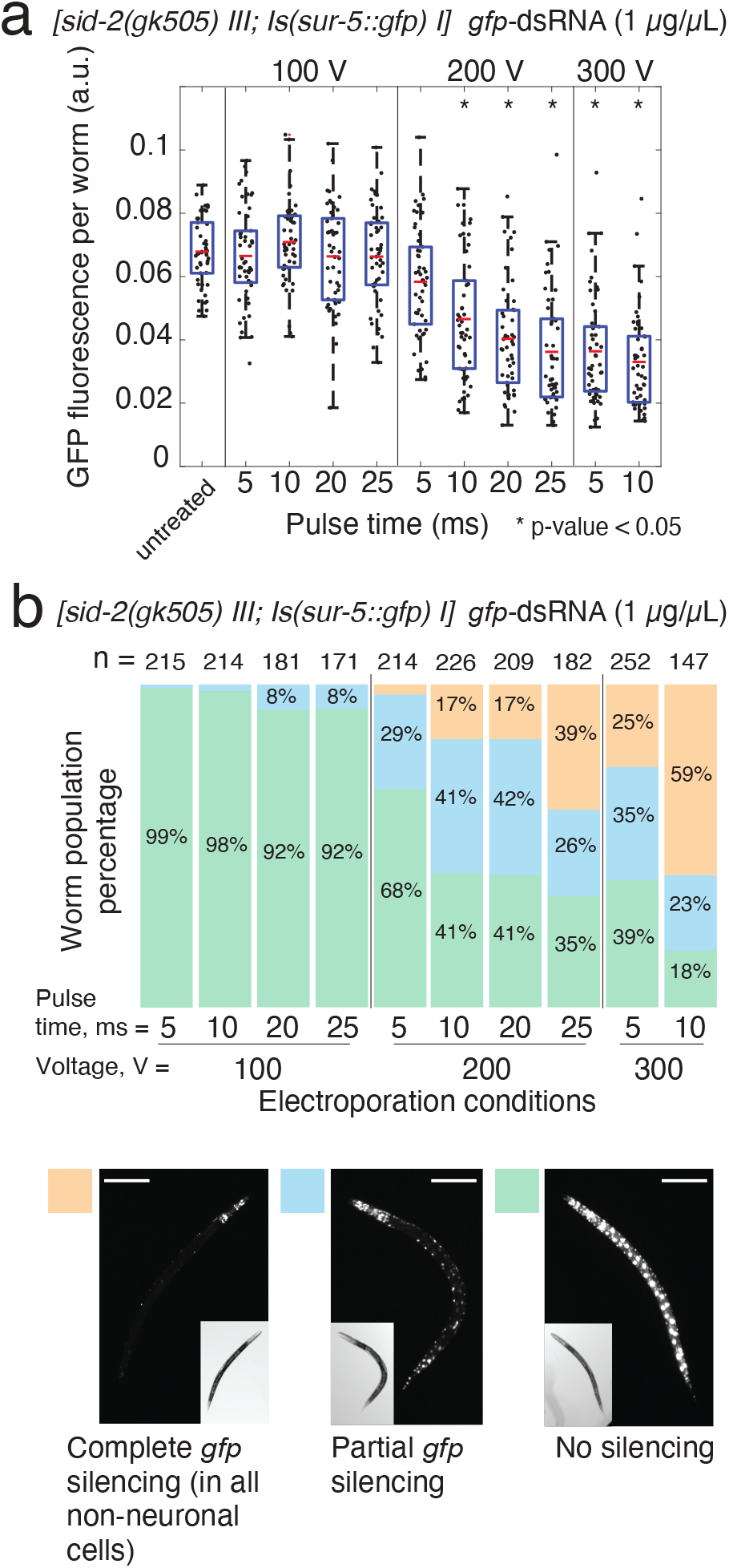
Identification of electroporation conditions for efficient delivery of dsRNA in *C.elegans*. To evaluate the effectiveness of nucleic acid delivery into animals, we used highly sensitive RNAi-mediated silencing of a GFP transgene following electroporation of dsRNA. **(a)**Synchronized L1 populations of BIG0107 *[sid-2(gk505) III;Is(sur-5:gfp) I]* worms (*~*250) were electroporated with *gfp*-dsRNA of 1 *μ*g/*μ*L using favorable electroporation conditions. Animals placed in electroporation buffer without dsRNA or electric discharge were used as “untreated” controls. For each condition, GFP fluorescence intensity of worms (n=50) was measured in arbitrary units (a.u.). Asterisk (*) indicates groups where significant *gfp* silencing compared to the untreated control was observed (p - value *<* 0.05, ANOVA test with Bonferroni correction). Red lines indicate means, blue boxes show 25^*th*^ and 75^*th*^ percentiles, whiskers show the data distribution range. **(b)**Three phenotypic categories of animals were scored in each condition group, including worms with “No silencing”, “Partial *gfp* silencing”, and “Complete *gfp* silencing” (all but neuronal cells).The electroporation parameters of 300 V 10 ms with the highest percentage in “Complete *gfp* silencing” category (59%) were chosen as the most efficient. n = number of worms scored. Representative images of worms from each category are shown, scale bar = 100 *μ*m.

### Determination of the tissue distribution of dsRNA delivery in *C.elegans*

With conditions for delivery optimized, we next aimed to identify the breadth of tissues that could be effectively electroporated. To test this, we utilized a similar reporter system together with the BIG0106 mutant defective in systemic RNAi, as SID-1 membrane channels facilitate spread of dsRNAs between tissues and into cells [Whangbo et al. 2017]. In this manner, *gfp* silencing should only be observed in those tissues and cells where *gfp*-dsRNA was directly delivered into the cell cytoplasm. Loss of systemic RNAi in these mutants predictably reduced the overall level of *gfp* silencing (**Figure 3**). Microscopic assessment of the animals indicated that silencing within hypodermal cells likely accounted for the majority of the significant decreases of GFP expression observed in *sid-1* mutants compared to controls (**Figure 3a-b, d**). These results do not exclude the possibility of delivery to other tissues, but suggest that the degree of delivery may be less efficient and would require additional optimization for silencing to occur. The presence of a large proportion of worms with partial *gfp* silencing (**Figure 2**) also suggests that the impact of the electric pulse along the animal body may not be uniform and depends on worm position in the cuvette that could lead to observed *gfp* silencing variations both between and within animals. Together these results indicate that electroporation delivers *gfp*-dsRNA most efficiently to hypodermal cells and then spreads to other tissues in a SID-1-dependent manner (**Figure 3c, e**).

**Figure 3:**
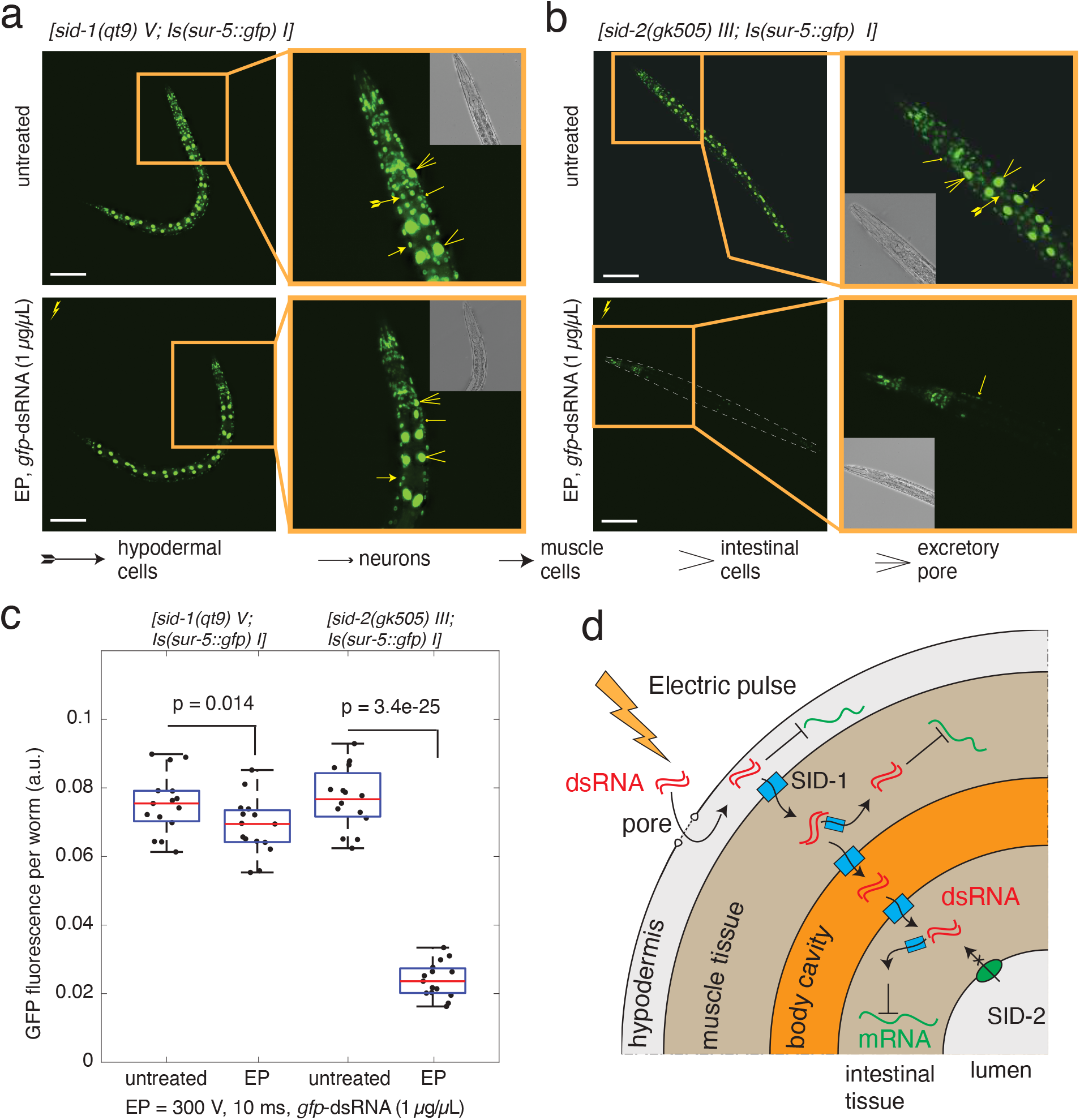
Evaluation of tissue distribution of RNAi silencing in electroporated animals. **(a, b)** Representative images of BIG0106 *[sid-1(qt9) V; Is(sur-5::gfp) I]* and BIG0107 *[sid-2(gk505) III; Is(sur-5::gfp) I]* worms were taken 48 hrs after the electroporation of L1 worm populations with *gfp*-dsRNA of 1 *μ*g/*μ*L using 300 V 10 ms conditions. Images of “untreated” control animals (no electroporation, no dsRNA) are presented for comparison. Scale bar = 100 *μ*m. **(c)** Levels of GFP fluorescence in both worm strains (n=15) after electroporation were compared to the untreated controls (P-values are noted, ANOVA test with Bonferroni correction). No significant differences in GFP fluorescence between untreated control worms from each strain were found. Red lines indicate means, blue boxes show 25^*th*^ and 75^*th*^ percentiles, whiskers show the data distribution range. **(d)** Schematic of the presumed routes of dsRNA transport highlighting hypodermal entry as a primary site of initial dsRNA delivery by electroporation, followed by spread to other tissues in a SID-1-dependent manner.

### Dose dependent delivery of dsRNA by electroporation

RNAi mediated silencing in *C.elegans* occurs in a dose dependent manner [Whangbo et al. 2017], which can be particularly useful when testing functions of essential genes. Since the experiments outlined above utilized highly concentrated levels of dsRNA (1 *μ*g/*μ*L), we next sought to identify whether we could control degree of silencing by titrating the levels of dsRNA targeting native genes delivered to the VC1119 animals. To test this, we selected two native genes expressed in the hypodermis with readily quantifiable size-based phenotypes, *nhr-23* (developmental arrest [Kouns et al. 2011]) and *dpy-13* (dumpy [von Mende et al. 1988]), to trigger silencing by different levels of dsRNA concentration (10 ng/*μ*L, 100 ng/*μ*L and 1 *μ*g/*μ*L). For each gene, we observed dose dependent, electroporation-driven ranges in silencing depending on the amount of dsRNA in solution (**Figure 4a,d**). Notably though, 100 fold less concentrated *nhr-23*-dsRNA was able to cause developmental arrests in 70% of animals compared to 96% for animals treated with 1 *μ*g/*μ*L of *nhr-23*-dsRNA (**Figure 4b-c**). While for *dpy-13*, we observed a lower penetrance of the dumpy phenotype and more gradual decrease in the average worm size with the increase of *dpy-13*-dsRNA concentration (**Figure 4d-f**). Together, these results illustrate that electroporation of dsRNA can titrate levels of gene silencing with minimal levels of starting dsRNA material.

**Figure 4:**
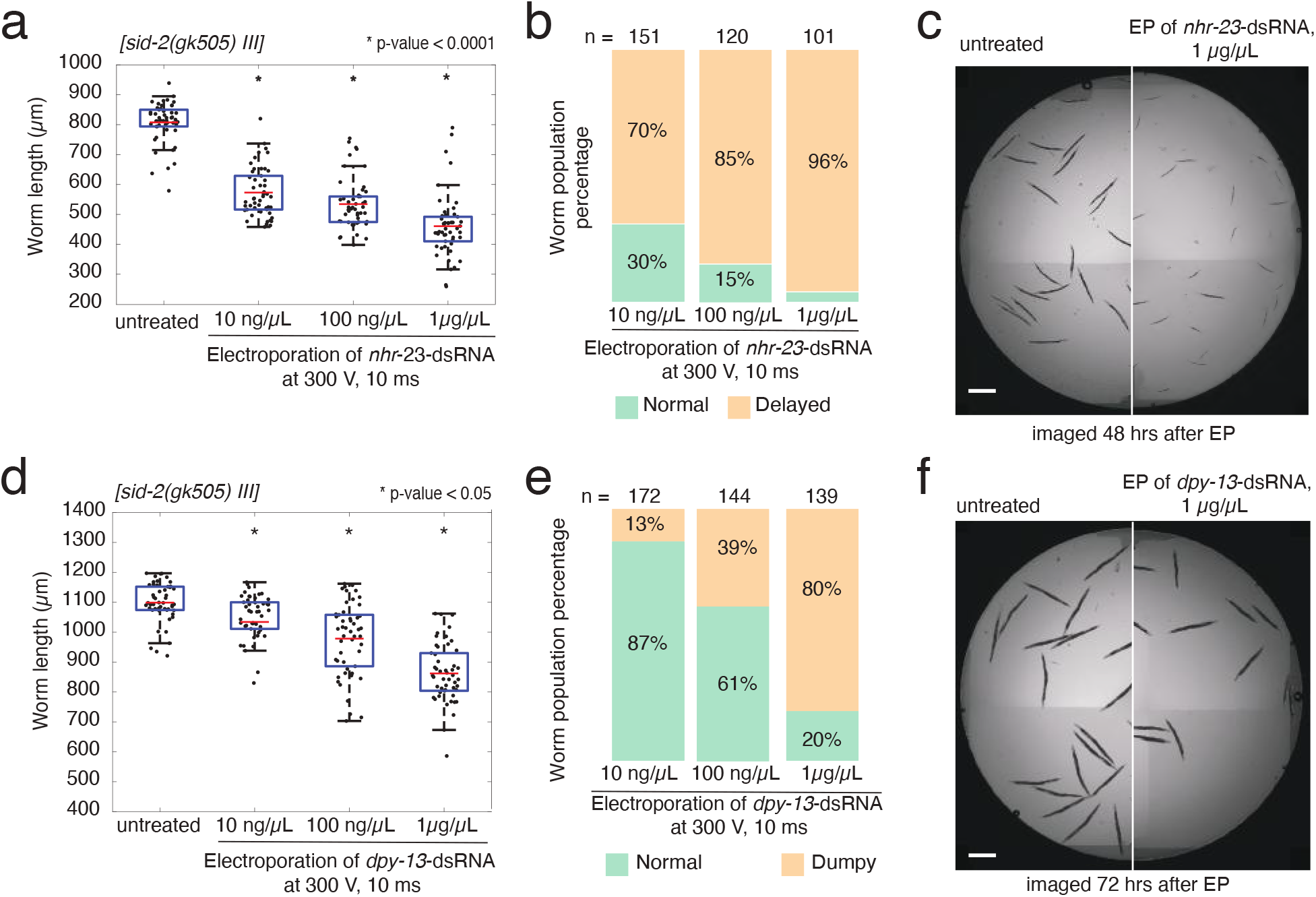
Efficiency of electroporation-driven gene silencing of endogenous genes is dose dependent. In order to test the effectiveness in non-transgenic animals, we targeted endogenous hypodermally expressed genes with robust RNAi phenotypes, *nhr-23* (larval arrest, **(a,b,c)**) and *dpy-13* (shortened body size, **(d,e,f)**). **(a)** Impact of electroporation of *nhr-23*-dsRNA on the development of *sid-2(gk505)* worms treated at L1 stage and imaged after 48 hrs. Red lines indicate means, blue boxes show 25^*th*^ and 75^*th*^ percentiles, whiskers show the data distribution range. **(b)** Proportion of animals scored as having either “normal” or “delayed” development after electroporation. **(c)** Representative image of worms show the *nhr-23* silencing effect at 1*μ*g/*μ*L of dsRNA (right image), when compared to untreated worms (left image). Scale bar = 500 *μ*m. **(d)** Impact of electroporation of *dpy-13*-dsRNA on body size of *sid-2(gk505)* worms treated at L1 stage and imaged after 72 hrs. Red lines indicate means, blue boxes show 25^*th*^ and 75^*th*^ percentiles, whiskers show the data distribution range. **(e)** Proportion of animals scored as “normal” or “dumpy” after electroporation. **(f)** Representative images of worms demonstrate the *dpy-13* silencing at 1 *μ*g/*μ*L of dsRNA (right image) in comparison with untreated control worms (left image). Scale bar = 500 *μ*m. Asterisk (*) indicates groups with significant gene silencing compared to the untreated control (p - value *<* 0.05, ANOVA test with Bonferroni correction).

### Germline delivery and transmission of electroporated dsRNA to progeny

Next we aimed to examine whether electroporation could be used to deliver material to the germline of animals. We expected the most efficient transmission of dsRNAs to occur in animals that are at or near reproductive maturity (i.e., L4 stage or older [Marra et al. 2016]). To test whether dsRNA can target the germline, populations of L4 animals VC1119 were electroporated with a germline-specific *pos-1*-dsRNA of 1 *μ*g/*μ*L (**Figure 5a**), as efficient silencing of the *pos-1* gene produces a robust embryonic lethal phenotype [Tabara et al. 1998]. After 24 hrs adult animals were removed from the plate and the progeny were scored for hatching after an additional 48 hrs. We observed consistent delivery and efficient *pos-1* silencing as evidenced by the prevalence of unhatched eggs from electroporated animals compared to those of untreated control animals (**Figure 5b**). This indicates that the hypodermally delivered dsRNA can spread and silence effectively in the germline, which we are not able to observe with *sur-5::gfp* strains due to intrinsic germline silencing of *gfp* transgenes. Additionally, we also tested *sid-1(qt9)* mutants defective in systemic RNAi and observed no difference in progeny development derived from electroporated population of worms compared to the control worms (**Figure 5b**). Together, these studies indicate transmission of electro-plated dsRNA to the germlines.

**Figure 5:**
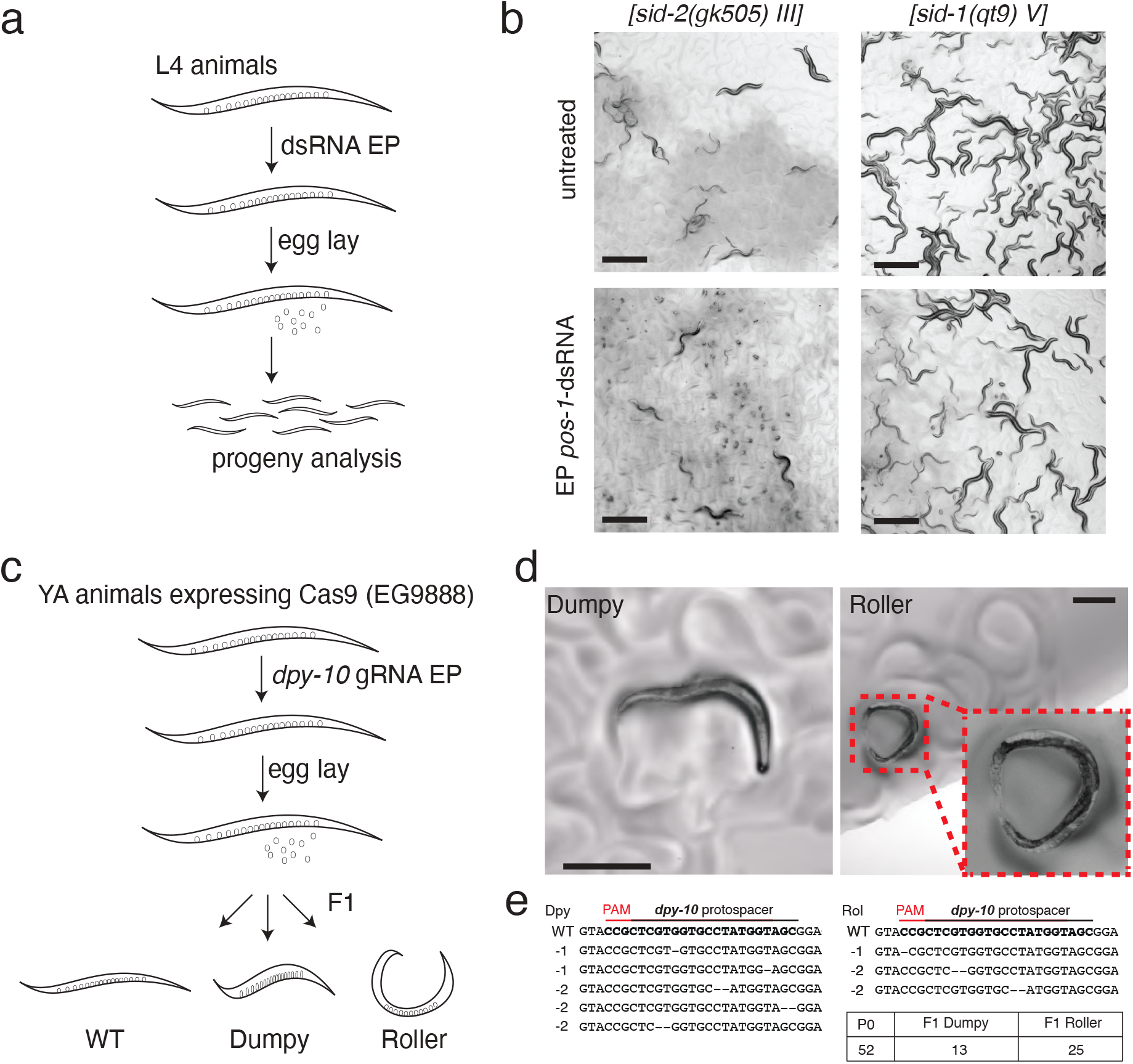
Evaluation of electroporation for delivery to the animal germline and progeny. To further test the utility of this approach, we sought to identify whether we could, first, stimulate RNAi knockdown of endogenous gene *pos-1* expressed in germline with robust phenotype (embryonic lethality) and, second, deliver guide RNA (gRNA) for CRISPR/Cas9-mediated genome editing of the endogenous *dpy-10* gene. **(a)** Schematic of *pos-1*-dsRNA delivery to L4 worms by electroporation (300 V for 10 ms, 1 *μ*g/*μ*L of dsRNA in electroporation buffer with final volume of 50 *μ*L) followed by phenotypic analysis of progeny. Animals were allowed to lay eggs for 24 hrs, removed from the lawn, and the proportion of hatched progeny was determined after 48 hrs. **(b)** Representative images of effective electroporation-mediated delivery of *pos-1*-dsRNA to *sid-2(gk505)* animals. Scale bar = 500 *μ*m. **(c)** Schematic of delivery of *dpy-10* gRNA to Young Adult (YA) animals by electroporation (300 V for 10 ms, 1 *μ*g/*μ*L of RNA in electroporation buffer with final volume of 50 *μ*L) followed by phenotypic screening of progeny for evidence of genome editing (Dumpy or Roller). **(d)** Representative images of successful electroporation of *dpy-10* gRNA in EG9888 animals that resulted in F1 progeny with visible Rol and Dpy phenotypes. Scale bar = 300 *μ*m. **(d)** Illumina sequencing-based confirmation of Cas9-mediated mutations.

### Evaluation of electroporation to deliver guide RNA to germlines for Cas9-mediated genome editing

Since we demonstrated that we could deliver dsRNAs to the germline, we sought to next determine whether that delivery could be extended to guide RNAs (gRNAs) for CRISPR/Cas9 based genome editing. Typically, gRNAs are injected along with additional components into the germlines of animals one-by-one to target disruption of specific genes [Prior et al. 2017]. In this study, we took advantage of transgenic worms stably expressing *cas9* in the germline (EG9888 *[W01A8.6(oxTi—-[Pmex-5::cas9(+ smu-2 introns), Phsp-16.41::Cre, Pmyo-2::2xNLS-CyOFP + lox2272])I]*; unpublished, a gift from Dr. Matthew Schwartz and Dr. Erik Jorgensen) that should only need introduction of gRNAs to facilitate targeting. We then chose to deliver a well characterized and robust gRNA targeting *dpy-10* that is commonly used as a co-CRISPR marker for CRISPR/Cas9 editing during microinjection [Arribere 2014]. In order to ensure robust Cas9 production, we electroporated *dpy-10*-gRNA (1 *μ*g/*μ*L; 300 V and 10 ms) into a population of young adult (YA) worms and screened F1 progeny for editing events both phenotypically and genetically (**Figure 5c**). As additional controls, we included both soaking in *dpy-10*-gRNA (1 *μ*g/*μ*L) and feeding on *E.coli* producing *dpy-10*-gRNA; neither of these controls produced phenotypically altered progeny. Electroporated *dpy-10*-gRNA was able to be successfully delivered in population of P0 worms (n = 52), and albeit at low levels resulted in F1 progeny production with Rol (n = 25) and Dpy (n = 13) phenotypes (**Figure 5d; File S2**). However, the observed phenotypic changes were not heritable or lethal and more likely were only somatic in F1 animals, as F2 progeny did not retain their phenotypes. Consistent with this notion, Illumina sequencing of F1 Rol and Dpy animals identified low indels frequency rates ranging from 0.2% - 1.3% with single and dinucleotide deletions (**Figure 5e**). Worth noting, in order to confirm functionality of *in-vitro* produced gRNA, EG9888 animals were injected with *dpy-10*-gRNA followed by F1 progeny selection with Rol and Dpy phenotypes. Subsequently, F2 progeny from these animals was also genotypically confirmed to inherit Rol and Dpy phenotypes (data not shown). Together, these results suggest that electroporation-based delivery of gRNAs is possible, but further optimization is needed to increase the efficiency of targeting moving forward.

## DISCUSSION

We demonstrate that nucleic acids can be delivered via electroporation into *C.elegans* worms at several stages of life. Electroporation conditions were optimized to maximize animal viability and potential for material delivery. Using RNAi as a sensitive readout for delivery of dsRNA, we show that electroporation-mediated delivery of *in-vitro* synthesized gene-specific dsRNAs resulted in RNAi silencing of both GFP-reporter transgenes and native genes, such as *nhr-23*, *dpy-13*, *pos-1*. Dose-dependent increase in electroporation-driven RNAi silencing was demonstrated with dsRNA concentrations ranging from 10 ng/*μ*L to 1 *μ*g/*μ*L. The use of *sid-1(qt9)* and *sid-2(gk505)* mutants with *sur-5::gfp* transgene reporter allowed us to dissect the way electroporated dsRNA enters inside the worm body. Namely, electroporated dsRNA is delivered into the cytoplasm of hypodermal cells and distributed systemically by SID-1 RNA channel throughout the body and into germlines (**Figure 3d**). The proposed electroporation method of population scale dsRNA delivery is quick, easy and can be accomplished in 15 minutes compared to traditional 24-48 hours needed for efficient RNAi by feeding and soaking [Ahringer 2006].

Being able to pair host genetic knockdowns that do not require alterations in the physiology of the animal are key to the usefulness of the system regardless of the question being interrogated. Studies of *C.elegans* commonly rely on standard *E.coli* OP50 diet in the laboratory and on RNAi screenings where the other *E.coli* strain HT115 is used both as a diet source and a producer of dsRNA. It was found that these two *E.coli* strains differentially affect gene expression profiles in worms [Coolon et al. 2009] and influence on animal metabolism, physiology, development, behavior, immunity, and lifespan. For this reason, recent advances have led to the development of an *E.coli* OP50 RNAi strain [Neve et al. 2020]. However, expanded appreciation for and widespread utilization of microbes from *C.elegans* natural microbiome [Zhang et al. 2017], each with their own impact on aspects of host physiology [Samuel et al. 2016], complicates this paradigm. Each strain would need its own RNAi library in order to properly examine the genetics of host-microbe interactions in these cases. Thus, we believe that electroporation as a bacteria-free dsRNA delivery method can mitigate the need to introduce another microbe into the mix (*E.coli*) in RNAi-based tests of host-microbe interactions. In addition, compared to RNAi silencing implemented via soaking, which is also a bacteria-free method, electroporation eliminates prolonged worm starvation or larval developmental arrest, which also has a pronounced effect on worm gene expression profiles particularly if completed early in life [Rechavi et al. 2014].

Beyond knockdowns, many effective strategies have been developed for precise genome editing of *C.elegans* [Dickinson and Goldstein 2016, Wang et al. 2018, Schwartz and Jorgensen, Au et al. 2019, Yang et al. 2020]. Nearly all of these strategies rely on low-throughput microinjection methods for delivery of nucleic acids mixtures. Here, we present proof-of-principle studies that electroporation may be a useful strategy for circumventing the microinjection step in these pipelines through population-scale delivery of guide RNAs in Cas9 expressing transgenic worms. We observed the Rol and Dpy phenotypes after electroporation of young adult worms with *dpy-10* gRNA only in the F1 generation, suggesting that phenotypes were presumably caused by editing in somatic cells. Further studies will be needed to determine whether this is due to the delivery route that the electroporated gRNA reached the germline, which is likely SID-1 dependent spreading from hypodermal cells in this case. Somatic editing in F1 generation of worms after microinjections of CRISPR/Cas9 complex in worm’s syncytial gonads is commonly observed and is a consequence of residual Cas9 activity in the fertilized embryos [Cho et al. 2013]. This explanation fits well to our experimental results given that in EG9888 transgenic strain Cas9 is expressed under cytoplasmic germline *mex-5* promoter which remains active in fertilized eggs as well [Tenlen et al. 2008]. Additionally, previous studies have demonstrated that transportation of dsRNA to proximal oocytes and embryos in mature worms also occurs via RME-2 mediated endocytosis [Marra et al. 2016]. It may be possible to engage this pathway for more efficient and timely transfer of gRNAs to the germline. Overall, we believe that our findings hold a promise for further development of population scale, electroporation-mediated delivery of nucleic acids into *C.elegans* for a wide variety of applications.

## ACKNOWLEDGEMENTS

This work was supported by an NIH New Innovator Award DP2DK116645 (to B.S.S) and pilot funding from the Dunn Foundation (to B.S.S.). Some strains were provided by the CGC, which is funded by NIH Office of Research Infrastructure Programs (P40 OD010440). We are grateful to the Gary Ruvkun, Craig Hunter and Erik Jorgensen laboratories for sharing *C.elegans* strains. We also thank members of the Robert Britton, Gretchen Diehl, David Reiner and Zheng Zhou laboratories for use of key equipment, support and advice, and members of the Samuel lab for helpful comments on the manuscript.

## SUPPLEMENTARY INFORMATION

### Supplementary data provided with the manuscript

**Table S1.** Strains used in this study.

| Strain name | Genotype | Source |
| --- | --- | --- |
| N2 |  | Samuel lab stock |
| HC196 | *[sid-1(qt9) V]* | CGC |
| VC1119 | *[dyf-2\&ZK520.2(gk505) III]*<br>(named as *[sid-2(gk505)III]* in text) | CGC |
| GR1403 | *[ls(sur-5::gfp) I; eri-1(mg366) IV]* | Gary Ruvkun lab |
| BIG0105 | *[ls(sur-5::gfp) I]* | this study; generated by crossing GR1403 and N2 |
| BIG0106 | *[sid-1(qt9) V; ls(sur-5::gfp) I]* | this study; generated by crossing BIG0105 and HC196 |
| BIG0107 | *[sid-2(gk505) III; ls(sur-5::gfp) I]* | this study; generated by crossing BIG0105 and VC1119 |
| EG9888 | *[W01A8.6(oxTi---{Pmex-5::cas9(+ smu-2 introns), Phsp-16.41::Cre, Pmyo-2::2xNLS-CyOFP + lox2272})I]* | Eric Jorgensen lab |

**Table S2.**
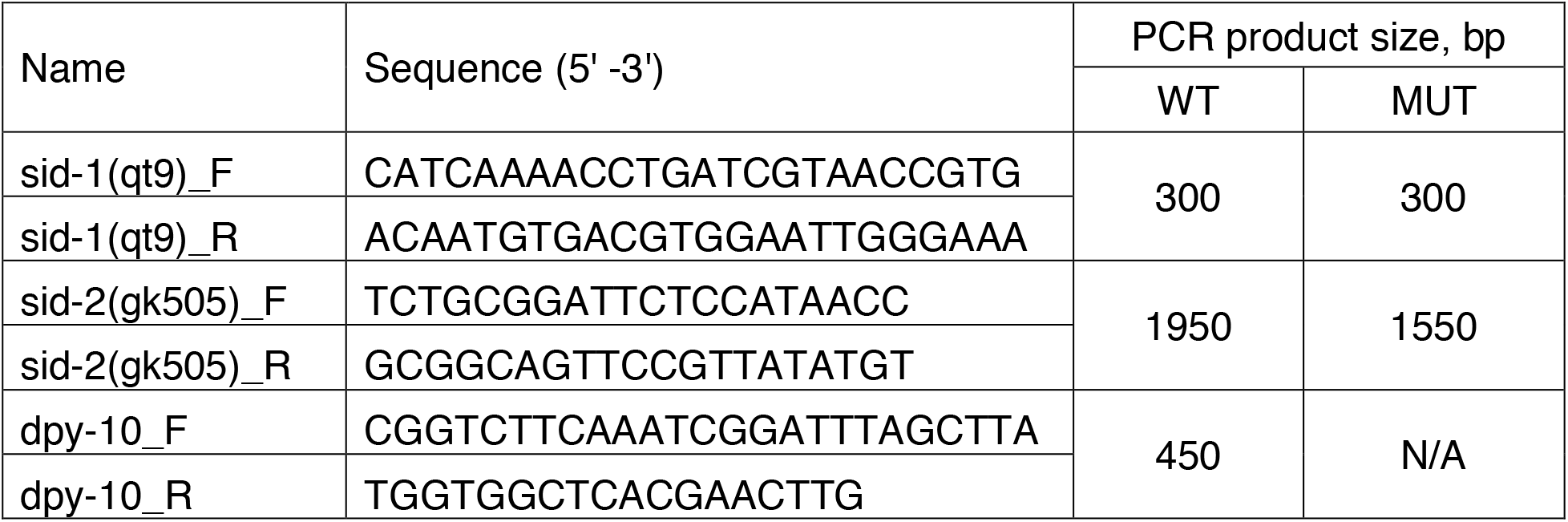
Primers used for genotyping of worms.

**Table S3.** Viability of worms (N2) after electroporation at L4 stage.

| **Voltage, V** | **Pulse time, ms** | **Initial # worms taken** | **# worms alive in 24 hours** | **Viability rate, %** |
| --- | --- | --- | --- | --- |
| **100** | **5** | 150 | 139 | 93 |
|  | **10** | 124 | 112 | 90 |
|  | **20** | 168 | 146 | 87 |
|  | **25** | 165 | 146 | 88 |
| **200** | **5** | 169 | 160 | 95 |
|  | **10** | 286 | 259 | 91 |
|  | **20** | 236 | 212 | 90 |
|  | **25** | 193 | 177 | 92 |
| **300** | **5** | 266 | 257 | 97 |
|  | **10** | 337 | 321 | 95 |
|  | **20** | 70 | 51 | 73 |
|  | **25** | 121 | 86 | 71 |
| **400** | **5** | 119 | 103 | 87 |
|  | **10** | 123 | 91 | 74 |
|  | **20** | 90 | 48 | 53 |
|  | **25** | 108 | 2 | 2 |
| **Untreated** |  | 205 | 175 | 85 |

### • File S1. Custom Matlab scripts for image analysis

The file contains two Matlab scripts for image analysis used throughout the research, additional files requared for the scripts (map.mat; bfmatlab folder) and exemplary fluorescent images (in .nd2 format) of “Control”, “Partially silenced” and “Completely silenced” populations of worms after electroporation with gfp-dsRNA.

#### – Categorical counting script

The script was used to count the number of worms belonging to the following three categories of worms in population: partial silencing, complete silencing, and no silencing. As an input the script accept .nd2 image files taken on Nikon fluorescent microscope. The file should have resolution of 6964 *×* 6964 pixels and two channels (bright field and fluorescence field). During script execution, the script provides user with hints on what to do during each stage. Briefly, on a bright field image first manually select worms by mouse clicking, when finished click once inside the square in left upper corner; next go on fluorescent image and select worms corresponding to one category, by clicking, when finished click once inside the square in left upper corner, then continue selection of worms from the other category, again when finished click once inside the square in left upper corner; at the end you will see the number of worm in each of three categories and a total number of worms. As an output the script generates a table with a number of worms falling into each category.

#### – Single channel worm selection script

The script was used for measuring average fluorescence intensity of the worm (or multiple worms) selected on the image. As an input the script accepts .nd2 image files taken on Nikon fluorescent microscope. The file should have resolution of 6964 *×* 6964 pixels and two channels (bright field and fluorescence field). During script execution, the script provides user with hints on what to do during each stage. The output file of the script is a table containing a number of rows corresponding to the selected worms. Each raw for each particular worm includes the following information: worm area (Column B), average fluorescence normalized to the worm’s area (Column C), average background intensity (Column F) and average fluorescence intensity with subtracted background intensity (Column H). Average background intensity is measured on area with no worm. The script also generates a separate folder where the images for bright field, fluorescent field and binary masks for each worm are saved.

### • File S2. Videos of live worms with Dpy and Rol phenotypes

The file contains three vide files (in .mp4 format). The video files demonstrate representative F1 worms with Dumpy and Roller phenotypes observed after electroporation of *dpy-10* gRNA in the population of YA EG9888 worms with Cas9 expression in the germline.

### • File S3. Illumina amplicon sequencing data of dpy-10 loci in Dpy and Rol worms

The file contains NGS sequencing files (in .fastq.gz format) including:

- Dumpy_R1.fastq.gz
- Dumpy_R2.fastq.gz
- Roller_R1.fastq.gz
- Roller_R2.fastq.gz

The files are Illumina 2×250 pair-end sequencing datasets of dpy-10 PCR amplicons obtained from the single worms with Dumpy and Roller phenotypes.

### • File S4. Table with raw data of worm body lengths and GFP fluorescence intensities measurements after electroporation with dsRNAs

The file contains additional information and raw data including:

- Table S4.1 (Sheet 1) - raw data Figure 1b; animals viability testing after electroporation at L1 stage under different electroporation conditions
- Table S4.2 (Sheet 2) - raw data for Figure 1c; measurement of lengths of worms after electroporation at different conditions
- Table S4.3 (Sheet 3) - raw data for Figure 2a; measurement of GFP fluorescence intensity per worm
- Table S4.4 (Sheet 4) - raw data for Figure 3c; measurement of GFP fluorescence intensity per worm
- Table S4.5 (Sheet 5) - raw data for Figure 4a; measurement of lengths of worms after electroporation at different dsRNA concentrations
- Table S4.6 (Sheet 6) - raw data for Figure 4d; measurement of lengths of worms after electroporation at different dsRNA concentrations.

